# Juvenile behavioral alterations and sex-dependent deficits in hippocampal neurogenesis following early-life stress in rats

**DOI:** 10.64898/2026.09.15.751844

**Authors:** Ailen Alba Colapietro, Jazmín Grillo Balboa, Karen Melany Stefani, Marianela Noemí Ceol Retamal, María Eugenia Pallarés, Marta Cristina Antonelli, Silvina Laura Diaz

## Abstract

Early-life adversity during critical developmental windows markedly increases vulnerability to neuropsychiatric disorders later in life. Although adult behavioral and neurobiological alterations following early stress are well documented, the early emerging phenotypes and their underlying cellular and molecular mechanisms during juvenile development remain poorly understood. In this study, we utilized the translational Scarcity-Adversity Model (SAM) to investigate early behavioral alterations and hippocampal neuroplasticity in juvenile offspring. We characterized the behavioral profile of juvenile subjects exposed to SAM to identify early manifestations of vulnerability. Next, we evaluated hippocampal neurogenesis by quantifying neural progenitor cell proliferation immediately following stress exposure (at postnatal day 13, PND 13) and the short-term survival of newborn neurons (1 week post-labeling) in the dentate gyrus of juvenile offspring. Furthermore, given the functional compartmentalization of the hippocampus, we measured the protein expression of mature BDNF (mBDNF) and its precursor proBDNF across both dorsal and ventral domains. At the behavioral level, SAM-exposed juvenile offspring exhibited increased immobility in the forced swim test alongside impaired spatial memory in the novel object location task. Regarding hippocampal neurogenesis, SAM exposure did not alter immediate progenitor proliferation at PND 13; however, it produced a significant, male-specific reduction in newborn neuron survival, concomitant with a trend toward decreased mBDNF levels in males. Together, these results provide critical insight into the early neurodevelopmental trajectories triggered by early-life adversity, identifying early behavioral impairments and male-specific neuroplastic deficits as key juvenile markers of vulnerability preceding adult psychopathology.

## INTRODUCTION

Exposure to chronic stress during critical developmental windows, including aberrant caregiver–infant interactions, markedly heightens susceptibility to neuropsychiatric disorders. During this sensitive period, primary caregivers provide vital scaffolding for stress buffering and emotional regulation, coinciding with the structural and functional maturation of circuits underlying socio-emotional attachment (Callaghan et al., 2019). Extensive clinical and experimental evidence demonstrates that early adversity induces long-lasting alterations in neural architecture, predisposing individuals to severe psychiatric comorbidities, particularly anxiety and depression (Agorastos et al., 2019; Felitti et al., 1998; Heim & Nemeroff, 2001). Today, these disorders represent a major global public health burden due to their high prevalence, multifaceted clinical manifestations - ranging from mood dysregulation to persistent cognitive deficits - and substantial socioeconomic impact. In this context, elucidating the factors that drive differential vulnerability and identifying their earliest manifestations are essential for developing timely, more effective therapeutic interventions (Condon et al., 2018).

To model these developmental dynamics in a controlled preclinical setting, the Scarcity-Adversity Model (SAM) has emerged as a robust translational paradigm to study early-life adversity (Raineki et al., 2010; Roth & Sullivan, 2005). Under this paradigm, we previously reported aberrant maternal behaviors, including rough handling, jumping over, and stepping on pups, alongside reduced nursing, which translated into elevated stress levels in the offspring (Colapietro et al., 2025). Importantly, the long-term consequences of this early adversity revealed distinct, sex-dependent behavioral alterations in adulthood (Grillo Balboa et al., 2025). Given that these neurobiological and behavioral phenotypes begin to emerge during the post-weaning juvenile period, dissecting the cellular and molecular substrates involved represents a critical step for early intervention.

In this context, the hippocampus stands out as a primary target of early-life stress. In humans, childhood physical and emotional neglect has been linked to reduced hippocampal volume in patients with major depressive disorder, an effect that is notably more pronounced in males (Frodl et al., 2010). At the cellular level, the subgranular zone of the dentate gyrus constitutes one of the primary neurogenic niches in the adult mammalian brain (Gage, 2000). Because adult hippocampal neurogenesis is critical for higher cognitive functions - such as learning, memory consolidation, and stress-response regulation - its dysregulation is closely associated with various psychiatric and neurodegenerative conditions (Apple et al., 2016; Wu & Zhang, 2023). Notably, beyond the initial proliferation of neural progenitor cells, the long-term survival and functional integration of newborn neurons represent a critical checkpoint that is particularly vulnerable to chronic stress (Kempermann et al., 2015).

Therefore, to determine whether emergent behavioral phenotypes serve as early functional readouts of hippocampal vulnerability following early-life adversity, this study evaluated the impact of SAM from behavior to neuroplasticity. We first characterized the juvenile behavioral repertoire across emotional and cognitive domains to identify early-onset phenotypes. Guided by these behavioral outcomes - particularly in spatial memory and stress-coping strategies - we next examined their underlying cellular and molecular correlates within the hippocampus. Specifically, we quantified acute neural progenitor proliferation (at PND 13) and 1-week newborn neuron survival (at PND 28) in the dentate gyrus, alongside the regional expression of mature BDNF and proBDNF along the longitudinal (dorsal/ventral) hippocampal axis.

## MATERIAL AND METHODS

### Animals

A total of 24 pregnant adult female Wistar rats (weighing 250–300 g) were provided by the animal breeding facility of the “Instituto de Biología Celular y Neurociencias (IBCN, UBA-CONICET)”. Gestation was confirmed by detecting a vaginal plug after mating alongside consistent maternal weight gain. Animals were maintained on a 12:12 h light/dark schedule (lights off at 19:00 h) at a regulated temperature (23 ± 2 °C) and humidity range (30–70%), with standard commercial rodent chow (Asociación de Cooperativas Argentinas, Buenos Aires, Argentina) and tap water available ad libitum. All experimental procedures were reviewed and approved by the local Institutional Animal Care and Use Committee (CICUAL, Facultad de Medicina, Universidad de Buenos Aires; Protocol RESCD-2023–869-E-UBADCT#FMED).

Animal housing, birth tracking, and litter standardization were performed as previously described (Colapietro et al., 2025). Briefly, pregnant dams were housed in groups until gestational day 19–20, then individually caged until delivery. The day of birth (postnatal day 0, PND 0) was designated according to established observation windows (W. Chen et al., 2014). On PND 2, litters were adjusted to 4–6 pups per sex per dam. Pups were weaned on PND 21 and housed by sex and experimental condition (3–5 per cage). Dams were randomly allocated to either the Control or SAM group (n = 12 per group) one week postpartum. Behavioral testing was conducted in 44 juvenile offspring between PND 23 and 29. Brain tissue for BDNF analysis was collected from pups at PND 13 (n = 26) and PND 28 (n = 24). Cell proliferation was assessed at PND 13 (n = 28), and the survival of newborn neurons was measured at PND 28 (n = 30). The remaining offspring were used for other experiments (Colapietro et al., 2025; Grillo Balboa et al., 2025).

### Scarcity-Adversity Model (SAM)

SAM was implemented as previously described (Moriceau et al., 2009; Raineki et al., 2012). Briefly, litters were exposed to resource scarcity from PND 8 to PND 12 to induce altered maternal care and nesting deficits. During this period, control dams were provided with standard nesting material, consisting of a 4 cm layer of wood shavings and a full sheet of cotton paper. In contrast, SAM dams received limited nesting resources, comprising less than 1 cm of wood shavings and one-quarter sheet of cotton paper. Outside this developmental window (prior to PND 8 and following PND 12), all cages were maintained under standard control conditions.

### Behavioral Assessment of Emotional and Cognitive States

Behavioral testing was conducted in offspring between PND 23 and 29, ordered sequentially from the least to the most stressful paradigm at a rate of one test per day. All sessions were digitally recorded under controlled lighting and noise conditions. Specific behavioral parameters were manually scored by an observer blind to experimental groups using Solomon Coder software (version beta 19.08.02). Additionally, total distance traveled in the Open Field Test were tracked and automatedly quantified using Kinovea software (version 0.8.27).

### Open-Field Test (OFT)

Exploratory and anxiety-like behaviors were assessed in a square black arena (65 × 65 cm, 47 cm high walls). Rats were individually placed in the center of the apparatus and allowed to freely explore for 5 min. At the end of each trial, animals were returned to their home cages and the arena was thoroughly cleaned with 70% ethanol to eliminate olfactory cues. Total distance traveled, time spent in the central area, and the number of entries into the center were quantified. Exploratory activity was further categorized into supported rearing (vertical exploration against the walls) and unsupported rearing (vertical exploration without wall contact), serving as measures of general locomotion and emotional or risk-assessment state, respectively (Sturman et al., 2018).

### Elevated Plus Maze (EPM)

Anxiety-like behavior was evaluated following previously described protocols (Pastor et al., 2018). The maze consisted of two open arms (45 × 10 cm) and two enclosed black arms (45 × 10 × 50 cm) extending from a central platform (10 × 10 cm), elevated 65 cm above the floor. Rats were placed individually at the central intersection facing an open arm and allowed to explore for 5 min. An arm entry was defined as the placement of all four paws into the respective arm. Time spent in the central platform, open arms, and closed arms was quantified. Total arm entries were recorded as a measure of overall locomotor activity. Anxiety levels were evaluated as the percentage of time spent in the open arms relative to the total time spent across all arms, as well as the percentage of entries into the open arms relative to the total number of arm entries.

### Light-Dark Box (LDB)

Anxiety-like responses were evaluated using a two-compartment apparatus consisting of an illuminated white compartment (31 × 30 × 30 cm; 400 lx) and an enclosed dark compartment (15 × 30 × 30 cm), connected by a floor-level opening (12 × 8 cm) allowing free transition between chambers. Each rat was placed in the center of the illuminated compartment facing away from the dark partition and allowed to explore both chambers for 5 min. Total time spent in each compartment and the number of transitions between compartments were recorded.

### Forced Swim Test (FST)

Behavioral despair and stress-coping strategies were assessed as previously described by Pallarés et al. (2021). Rats were placed individually into a transparent glass cylinder (60 cm in height, 20 cm in diameter) filled with tap water (23– 25 °C) to a depth of 30 cm, preventing animals from supporting themselves on the bottom. On Day 1, rats underwent a 15-min habituation session. Twenty-four hours later, animals underwent a 5-min test session, which was recorded for behavioral analysis. The duration of the following behaviors was scored: (i) swimming, defined as active movements of the forepaws propelling the animal across the cylinder surface; (ii) climbing, defined as vigorous upward movements of the forepaws against the cylinder walls; and (iii) immobility, defined as passive floating with only the minimal movements necessary to keep the head above water. Immediately after each session, rats were dried with paper towels and returned to their home cages.

### Splash Test

Motivational state and self-care behavior were evaluated using the splash test according to Roversi et al. (2019). Rats were placed individually into a standard cage covered with a wire lid to ensure uninterrupted observation. A 10% sucrose solution was squirted onto the dorsal coat of the animal, and grooming activity was recorded over a 5-min period. Total grooming duration and frequency were scored, including nose and face grooming, head washing, and body cleaning.

### Object Location Task (OLT)

Spatial memory was assessed as described by Hattiangady et al. (2014) in an open-field arena (100 × 100 × 60 cm). The protocol comprised three 5-min phases separated by 60-min inter-trial intervals: a habituation phase with free exploration of the empty arena, a sample phase with exposure to two identical objects placed in opposite quadrants, and a test phase. During the test phase, objects were identical to those used in the sample phase, but one was relocated to a novel spatial position. The identity and position of the displaced object were counterbalanced across animals and litters to control for side bias. Object exploration was defined as directing the nose toward the object at a distance equal to or less than 2 cm, excluding climbing onto the object. The apparatus and objects were cleaned with 70% ethanol between trials. The Place Discrimination Index was calculated as the time spent exploring the object in the novel location divided by the total time spent exploring both objects, expressed as a percentage.

### Novel Object Recognition (NOR) Task

Recognition memory was evaluated in the same arena following an identical three-phase paradigm (habituation, sample, and test; 5 min each, 60-min inter-trial interval). During the test phase, one familiar object was replaced with a novel object of distinct shape and texture. Object exploration criteria and cleaning procedures were identical to those described for the OLT. The Object Discrimination Index was calculated as the time spent exploring the novel object divided by the total time spent exploring both objects, expressed as a percentage.

### Hippocampal Cell Proliferation and Survival Assays

#### Tissue Collection and Processing

To evaluate hippocampal cell proliferation, an independent cohort of pups was sacrificed at postnatal day (PND) 13. For cell survival assays, pups at PND 21 were weighed and received two intraperitoneal injections of the thymidine analogue 5-bromo-2′-deoxyuridine (BrdU; Sigma-Aldrich, B9285; 50 mg/kg each, administered 2 h apart; dissolved in 0.9% NaCl at 50 °C) and were maintained until PND 28 (1 week neuronal survival).

At the designated time points (PND 13 or PND 28), animals were deeply anesthetized with a ketamine/xylazine mixture (70 and 10 mg/kg, respectively) and transcardially perfused with heparinized saline (0.9% NaCl), followed by 4% paraformaldehyde in 0.1 M phosphate buffer (pH 7.4). Brains were dissected, post-fixed in the same fixative for 24 h at 4 °C, and cryoprotected in a 30% sucrose solution.

Serial 50-µm-thick coronal brain sections encompassing the entire rostro-caudal extent of the hippocampus were obtained using a freezing microtome and collected into six parallel series (yielding a 300-µm inter-section interval and an average of 13 sections per series). Sections containing the lateral ventricles were included as technical positive controls due to their high baseline cell proliferation rate. Free-floating sections were stored at −20 °C in a cryoprotectant solution (30% glycerol and 30% ethylene glycol in 0.1 M PBS) until processing.

#### Immunofluorescence

- Ki67 Detection (PND 13): To evaluate cell proliferation, free-floating brain sections from PND 13 animals were processed for Ki67 immunofluorescence. Sections were permeabilized and blocked against non-specific binding with 0.2% gelatin and 0.5% Triton X-100 in PBS for 1 h at room temperature (RT). Sections were then incubated overnight at 4 °C with a rabbit anti-Ki67 primary antibody (1:1000; Millipore, Cat# AB9260). Following PBS washes, sections were incubated for 2 h at RT with a goat anti-rabbit Alexa Fluor 568-conjugated secondary antibody (1:1000; Invitrogen, Cat# A11011/A11036), and cell nuclei were counterstained with Hoechst 33342 (1:10,000; Invitrogen, Cat# H3570 / 33342) for 10 min.

- BrdU Detection (PND 28): To visualize surviving newborn cells, free-floating sections from PND 28 animals were processed for BrdU immunofluorescence. For antigen retrieval, DNA denaturation was performed by incubating sections in 2 N HCl for 1 h at RT. Sections were then permeabilized and blocked with 0.2% gelatin and 0.5% Triton X-100 in PBS for 1 h at RT. Subsequently, sections were incubated overnight at 4 °C with a mouse anti-BrdU primary antibody (1:1000; Developmental Studies Hybridoma Bank, concentrate). After extensive PBS washes, sections were incubated for 2 h at RT with a goat anti-mouse Alexa Fluor 488-conjugated secondary antibody (1:1000; Invitrogen, Cat# A11029). Cell nuclei were counterstained with Hoechst 33342 (1:10,000; Invitrogen, Cat# 33342) for 10 min.

### Cell Quantification and Volumetric Analysis

Quantification of labeled cells (BrdU or Ki67) was conducted using an inverted epifluorescence microscope (Olympus IX81) under a 20× objective. In the case of Ki67, cell counts were restricted to the subgranular zone (SGZ), operationally defined as a two-cell-body-wide band along the inner margin of the dentate gyrus (DG) granule cell layer. BrdU+ cells were detected all along the granular zone. Cells exhibiting complete and homogeneous nuclear labeling were scored as positive.

To determine cell density, the area of the dorsal and ventral DG was estimated from 4× epifluorescence micrographs by delineating the granule cell layer area using ImageJ software (NIH). Cell density was calculated as the total estimated number of labeled cells divided by the total estimated area for each respective region.

### Western Blot Analysis of proBDNF and mBDNF Protein Levels

Tissue samples were homogenized in 250 µL of ice-cold RIPA buffer (50 mM Tris-HCl, 150 mM NaCl, 1% NP-40, 0.5% sodium deoxycholate, 0.1% SDS) supplemented with protease inhibitors (0.1 µM aprotinin, 0.1 µM pepstatin, 0.1 µM leupeptin, 0.2 mM PMSF) and centrifuged at 13,000 rpm for 30 min at 4 °C. Supernatants were collected, and total protein concentration was determined using the Bradford assay.

Protein samples (100 µg per lane mixed with 5× Laemmli loading buffer) were resolved by SDS-PAGE on 15% polyacrylamide gels and electrotransferred onto nitrocellulose membranes using the Mini-PROTEAN Tetra System (Bio-Rad) for 1 h. Membranes were blocked with 5% non-fat dry milk in Tris-buffered saline containing 0.1% Tween-20 (TBST) for 1 h at RT and subsequently incubated overnight at 4 °C with the following primary antibodies: mouse anti-BDNF (1:2000; Icosagen), mouse anti-proBDNF (1:2500; GeneCopoeia), and mouse anti-β-actin (1:1000; Santa Cruz Biotechnology) as an internal loading control.

After washing with TBST, membranes were incubated with horseradish peroxidase (HRP)-conjugated secondary antibodies (1:10,000; Jackson ImmunoResearch) prepared in blocking solution for 2 h at RT. Immunoreactive bands were visualized by enhanced chemiluminescence using the ECL Plus Western Blotting Substrate (Thermo Fisher Scientific) and captured on a GeneGnome imaging system (Syngene). Densitometric analysis was conducted using ImageJ software (NIH). For quantification, background-subtracted optical density values for mature BDNF (mBDNF, ∼15 kDa) and proBDNF (∼32 kDa) were normalized to the corresponding β-actin signal (42 kDa). Protein levels are expressed as relative optical density values.

### Statistical Analysis

All statistical analyses were performed using R software (v4.3.0; R Core Team, 2023) and RStudio (v2024.4.2.0), while graphical representations were generated using GraphPad Prism (v8.0.2). Data distribution normality and homoscedasticity were assessed using Q-Q plots, the Shapiro-Wilk test, standardized versus fitted residual plots, and Levene’s test.

For normally distributed data with homogeneous variances, linear models were fitted using two-way analysis of variance (ANOVA) with experimental group and sex as independent fixed factors. When significant main effects or interaction terms were detected, post hoc pairwise comparisons were conducted using Tukey’s honest significant difference (HSD) test via the emmeans package. For Western blot densitometric data, linear mixed-effects models (LMM; lme4 package v1.1-35.1) were applied, incorporating experimental group and sex as fixed factors and membrane ID (blot) as a random effect to account for inter-blot variability. Non-normally distributed datasets were analyzed using the non-parametric Kruskal-Wallis test followed by appropriate non-parametric pairwise comparisons. Statistical significance was set at p < 0.05. Significance levels for p-values are indicated as follows: *** p < 0.001, ** p < 0.01, * p < 0.05, and # p < 0.10 (statistical trend / approaching significance). Data are presented as mean ± SEM (or median with interquartile range for non-parametric datasets).

## RESULTS

### Juvenile SAM offspring exhibit passive stress-coping and spatial memory deficits without overt anxiety-like behaviors

We first evaluated the behavioral profile of juvenile offspring across a comprehensive test battery (Statistical analyses in Table 1). In the open field test, SAM offspring showed no significant differences in time spent in the center (Fig. 1.A), entries into the center, or total distance traveled compared to controls. In the elevated plus maze, open-arm exploration was unaffected, as shown by similar time spent (Fig. 1.B) and entries into the open arms across groups. In the light-dark box, the time spent in the illuminated compartment did not differ among groups (Fig. 1.C).

**Table 1.** Summary of statistical analyses for behavioral, cellular, and molecular assessments. Comprehensive breakdown of statistical evaluations across experimental paradigms and assays. For each measured parameter, the specific statistical test, planned comparisons, test statistics, degrees of freedom, associated p-values, and corresponding figure panel references are detailed. Significance codes for the p-value: ‘***’ <0.001, ‘**’ <0.01, ‘*’ <0.05, ‘#’ p < 0.10 (trends).

| Paradigm or essay | Measured parameter | Statistical test | Comparisons | Statistical | Degrees of freedom | p value | Fig |
| --- | --- | --- | --- | --- | --- | --- | --- |
| **Open-field test** | Time in the center | Kruskal-Wallis test | Exp. group | H=1.2315 | 1 | 0.2671 | 1 A |
|  |  |  | Sex | H=0.21296 | 1 | 0.6445 |  |
|  | Entrances to the center | 2-way ANOVA | Exp. group | F=0.9195 | 1 | 0.3430 |  |
|  |  |  | Sex | F=0.9195 | 1 | 0.3430 |  |
|  | Unsupported rearing time | 2-way ANOVA | Exp. group | F=3.4985 | 1 | 0.06824 # |  |
|  |  |  | Sex | F=1.2323 | 1 | 0.27312 |  |
|  | Supported rearing time | 2-way ANOVA | Exp. group | F=0.4935 | 1 | 0.4861 |  |
|  |  |  | Sex | F=1.5947 | 1 | 0.2133 |  |
|  | Distance traveled | 2-way ANOVA | Exp. group | F=0.1722 | 1 | 0.68027 |  |
|  |  |  | Sex | F=3.6694 | 1 | 0.06208<br># |  |
| **Elevated plus maze** | Time in open arms | Kruskal-Wallis test | Exp. group | H=2.4006 | 1 | 0.1213 | 1 B |
|  |  |  | Sex | H=0.09178 | 1 | 0.7619 |  |
|  | N° of entries to open arms | 2-way ANOVA | Exp. group | F=2.2059 | 1 | 0.1436 |  |
|  |  |  | Sex | F=0.1988 | 1 | 0.6576 |  |
| **Light-dark box** | Time in light compartment | Kruskal-Wallis test | Exp. group | H=0.85352 | 1 | 0.3556 | 1 C |
|  |  |  | Sex | H=0.36523 | 1 | 0.5456 |  |
|  | No. crossings between compartments | Kruskal-Wallis test | Exp. group | H= 2.669 | 1 | 0.1023 |  |
|  |  |  | Sex | H=0.50118 | 1 | 0.479 |  |
|  |  | Bonferroni | Control-SAM Males | H=2.133169 | 1 | 0.0987 # |  |
|  |  |  | Control-SAM Females | H=0.220950 | 1 | 1.0000 |  |
| **Forced swimming test** | Time spent immobile | 2-way ANOVA | Exp. group | F=31.5580 | 1 | **1.512e-06 \*\*\*** | 1 D |
|  |  |  | Sex | F=2.2583 | 1 | 0.1406 |  |
|  | Time spent climbing | Kruskal-Wallis test | Exp. group | H=5.2915 | 1 | **0.02143 \*** |  |
|  |  |  | Sex | H=1.0667 | 1 | 0.3017 |  |
|  | Time spent swimming | Kruskal-Wallis test | Exp. group | H=22.834 | 1 | **1.767e-06 \*\*\*** |  |
|  |  |  | Sex | H=0.081727 | 1 | 0.775 |  |
| **Splash test** | Time spent grooming | 2-way ANOVA | Exp. group | F=0.0861 | 1 | 0.7704 | 1 E |
|  |  |  | Sex | F=0.4085 | 1 | 0.5257 |  |
|  | Number of groomings | 2-way ANOVA | Exp. group | F=0.0386 | 1 | 0.8451 |  |
|  |  |  | Sex | F=0.6502 | 1 | 0.4239 |  |
| **Object location task** | Place discrimination index | 2-way ANOVA | Exp. group | F=4.2225 | 1 | **0.04573 \*** | 1 F |
|  |  |  | Sex | F=0.0874 | 1 | 0.76891 |  |
| **Novel object recognition** | Object discrimination index | 2-way ANOVA | Exp. group | F=0.6022 | 1 | 0.4415 | 1 G |
|  |  |  | Sex | F=2.1975 | 1 | 0.1446 |  |
| **Neuronal GD Proliferation** | Ki67<sup>+</sup> DG Dorsal Density (n°/mm<sup>2</sup>) | 2-way ANOVA | Exp. group | F=0.1063 | 1 | 0.74443 | 2 A |
|  |  |  | Sex | F=0.0001 | 1 | 0.99411 |  |
|  |  |  | Interaction | F=4.4159 | 1 | 0.03561<br>* |  |
|  |  | Tukey post hoc test | Control-SAM Males | T=-1.272 | 1 | 0.2246 |  |
|  |  |  | Control-SAM Females | T=0.666 | 1 | 0.5169 |  |
|  | Ki67<sup>+</sup> DG Ventral Density (n°/mm<sup>2</sup>) | Kruskal-Wallis test | Exp. group | H=0.60283 | 1 | 0.4375 |  |
|  |  |  | Sex | H=0.39796 | 1 | 0.5281 |  |
|  | Dorsal DG area (mm<sup>2</sup>) | 2-way ANOVA | Exp. group | F=0.7252 | 1 | 0.3944 | 2 B |
|  |  |  | Sex | F=0.3313 | 1 | 0.5649 |  |
|  |  |  | Interaction | F=0.0502 | 1 | 0.8227 |  |
|  | Ventral DG area (mm<sup>2</sup>) | 2-way ANOVA | Exp. group | F=0.4732 | 1 | 0.4915 |  |
|  |  |  | Sex | F=0.2247 | 1 | 0.6355 |  |
|  |  |  | Interaction | F=0.4916 | 1 | 0.4832 |  |
| **1 week neuronal survival** | BrdU<sup>+</sup> DG Dorsal Density (n°/mm<sup>2</sup>) | 2-way ANOVA | Exp. group | F=3.7747 | 1 | 0.06294<br># | 3 A |
|  |  |  | Sex | F=1.2082 | 1 | 0.28177 |  |
|  |  |  | Interactio<br>n | F=2.0091 | 1 | 0.16823 |  |
|  |  | Tukey post<br>hoc test | Control-SAM<br>Males | T=2.376 | 1 | **0.0252 \*** |  |
|  |  |  | Control-SAM<br>Females | T=0.372 | 1 | 0.7132 |  |
|  | BrdU<sup>+</sup> DG<br>Ventral<br>Density<br>(n°/mm<sup>2</sup>) | 2-way<br>ANOVA | Exp.<br>group | F=4.7744 | 1 | **0.03809**<br>* |  |
|  |  |  | Sex | F=<br>0.0089 | 1 | 0.92542 |  |
|  |  |  | Interactio<br>n | F=4.3801 | 1 | 0.04627<br>* |  |
|  |  | Tukey post<br>hoc test | Control-SAM<br>Males | T=3.025 | 1 | **0.0055**<br>** |  |
|  |  |  | Control-SAM<br>Females | T=0.065 | 1 | 0.9485 |  |
|  | Dorsal DG<br>area (mm<sup>2</sup>) | 2-way<br>ANOVA | Exp.<br>group | F=0.3408 | 1 | 0.5644 | 3 B |
|  |  |  | Sex | F=0.1827 | 1 | 0.6726 |  |
|  |  |  | Interactio<br>n | F=<br>2.8240 | 1 | 0.1048 |  |
|  |  | Tukey post<br>hoc test | Control-SAM<br>Males | T=1.601 | 1 | 0.1214 |  |
|  |  |  | Control-SAM Females | T=-0.775 | 1 | 0.4451 |  |
|  | Ventral DG area (mm<sup>2</sup>) | 2-way ANOVA | Exp. group | F=1.7786 | 1 | 0.1944 |  |
|  |  |  | Sex | F=0.3056 | 1 | 0.5853 |  |
|  |  |  | Interaction | F=2.2510 | 1 | 0.1461 |  |
|  |  | Tukey post hoc test | Control-SAM Males | T=2.002 | 1 | **0.0563 #** |  |
|  |  |  | Control-SAM Females | T=-0.150 | 1 | 0.8818 |  |
| **BDNF protein levels** | Dorsal mBDNF HPC protein levels (PND 13) | 2-way ANOVA | Exp. group | F=0.4412 | 1 | 0.50655 | 2 D |
|  |  |  | Sex | F=0.4607 | 1 | 0.49730 |  |
|  |  |  | Interaction | F=3.1380 | 1 | 0.07649 # |  |
|  |  | Tukey post hoc test | Control-SAM Males | T=1.752 | 1 | **0.0988 #** |  |
|  |  |  | Control-SAM Females | T=-0.673 | 1 | 0.5105 |  |
|  | Ventral mBDNF HPC protein levels | 2-way ANOVA | Exp. group | F=3.4737 | 1 | 0.06235 # |  |
|  | (PND 13) |  |  |  |  |  |  |
|  |  |  | Sex | F=0.6765 | 1 | 0.41078 |  |
|  |  |  | Interaction | F=4.9051 | 1 | 0.02678<br>* |  |
|  |  | Tukey post hoc test | Control-SAM Males | T=0.320 | 1 | 0.7534 |  |
|  |  |  | Control-SAM Females | T=-2.673 | 1 | **0.0177 \*** |  |
|  | Dorsal proBDNF HPC protein levels (PND 13) | 2-way ANOVA | Exp. group | F=0.1002 | 1 | 0.75163 | 2 E |
|  |  |  | Sex | F=3.7949 | 1 | 0.05141<br># |  |
|  |  |  | Interaction | F=0.7874 | 1 | 0.37489 |  |
|  | Ventral proBDNF HPC protein levels (PND 13) | 2-way ANOVA | Exp. group | F=0.1975 | 1 | 0.6567 |  |
|  |  |  | Sex | F=0.0087 | 1 | 0.9256 |  |
|  |  |  | Interaction | F=0.4774 | 1 | 0.4896 |  |
|  | Dorsal mBDNF HPC protein levels | 2-way ANOVA | Exp. Group | F=0.0120 | 1 | 0.91266 | 3 D |
|  | (PND 28) |  |  |  |  |  |  |
|  |  |  | Sex | F=4.5072 | 1 | 0.03375<br>* |  |
|  |  |  | Interaction | F=0.0387 | 1 | 0.84409 |  |
|  | Ventral<br>mBDNF<br>HPC protein<br>levels (PND<br>28) | 2-way<br>ANOVA | Exp.<br>group | F=0.6921 | 1 | 0.40545 |  |
|  |  |  | Sex | F=5.4044 | 1 | 0.02009<br>* |  |
|  |  |  | Interaction | F=0.0178 | 1 | 0.89396 |  |
|  | Dorsal<br>proBDNF<br>HPC protein<br>levels (PND<br>28) | 2-way<br>ANOVA | Exp.<br>group | F=0.0078 | 1 | 0.92973 | 3 E |
|  |  |  | Sex | F=0.0793 | 1 | 0.77819 |  |
|  |  |  | Interaction | F=3.2420 | 1 | 0.07177<br># |  |
|  |  | Tukey post<br>hoc test | Control-SAM<br>Males | T=-1.303 | 1 | 0.2098 |  |
|  |  |  | Control-SAM<br>Females | T=1.244 | 1 | 0.2302 |  |
|  | Ventral<br>proBDNF | 2-way<br>ANOVA | Exp.<br>group | F=0.3189 | 1 | 0.5723 |  |
|  | HPC protein<br>levels (PND<br>28) |  |  |  |  |  |  |
|  |  |  | Sex | F=0.1926 | 1 | 0.6608 |  |
|  |  |  | Interactio<br>n | F=1.1876 | 1 | 0.2758 |  |

**Figure 1.**
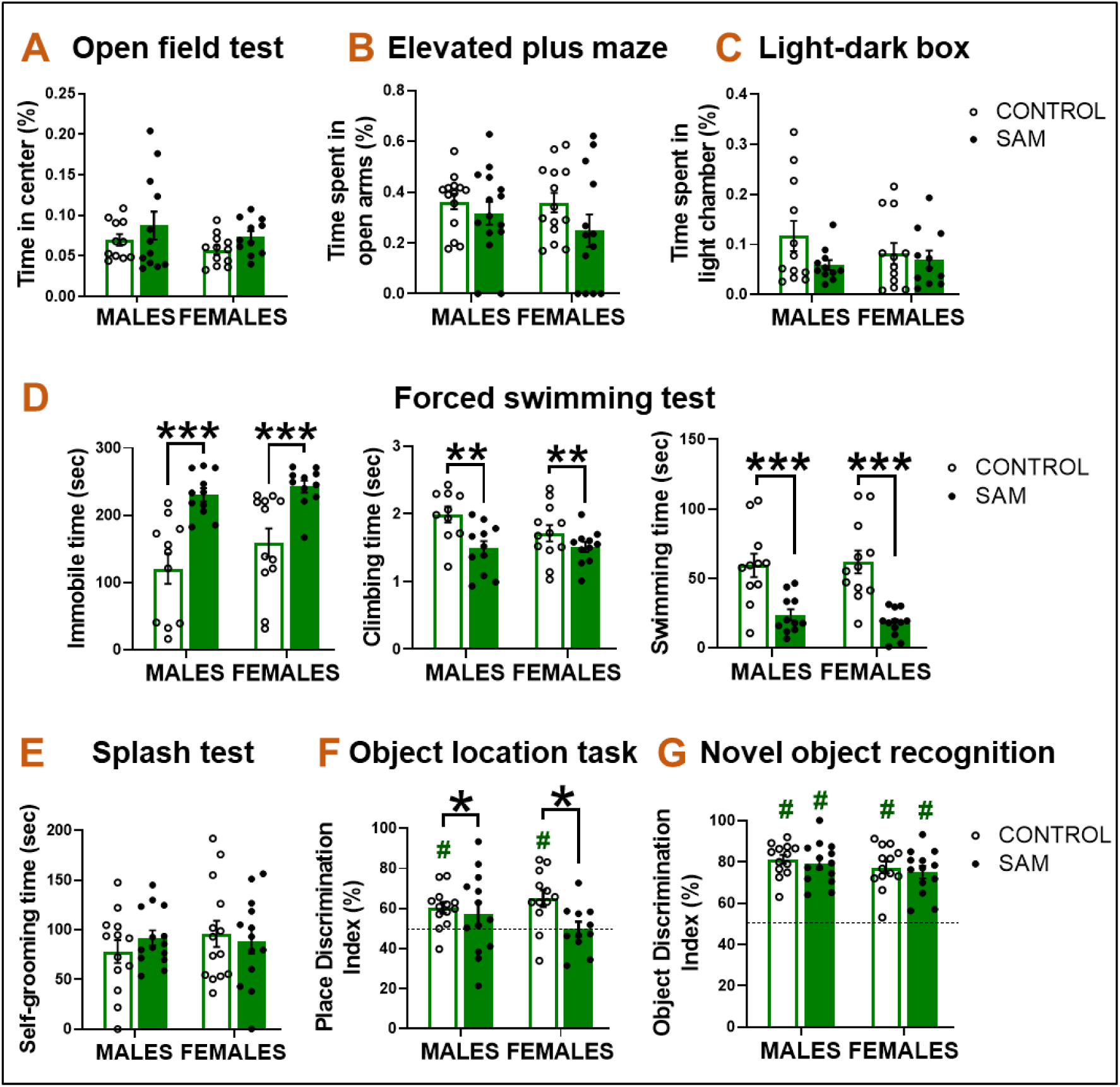
Behavioral profiling of juvenile offspring following early-life adversity. (A) Percentage of time spent in the center zone in the open field test. (B) Percentage of time spent in the open arms in the elevated plus maze. (C) Percentage of time spent in the light compartment in the light-dark box test. (D) Forced swim test: quantification of immobility, climbing, and swimming time. (E) Total grooming duration in the splash test. (F) Discrimination index in the object location test (OLT); the dashed line at 0.5 (50%) indicates chance level. (G) Discrimination index in the novel object recognition test (NOR). Data are presented as mean ± SEM (n = 8-10 animals per group). Two-way ANOVA followed by post-hoc test. # p < 0.05 vs. chance level (50%); * p < 0.05, ** p < 0.01, *** p < 0.001.

In the forced swim test, SAM offspring of both sexes exhibited a significant increase in immobility time, accompanied by a significant reduction in climbing and swimming behaviors relative to controls (Fig. 1.D). In the splash test, no significant differences were observed in grooming frequency (Fig. 1.E) or total grooming duration between groups. Cognitive assessment showed a significant reduced discrimination index, in the object location test (OLT), with SAM offspring of both sexes performing at chance level (no significant difference from 50%), demonstrating impaired spatial memory (Fig. 1.F). Conversely, intact recognition memory in the novel object recognition test (NOR) was registered, with SAM offspring displaying a discrimination index comparable to controls (Fig. 1.G).

### SAM alters regional mBDNF levels and subsequent neuronal survival without affecting immediate cell proliferation

Given that juvenile SAM offspring exhibited pronounced deficits in hippocampal-dependent spatial memory (OLT) and passive coping behavior, we next investigated whether these functional alterations were underpinned by disruptions in hippocampal neuroplasticity and neurotrophin signaling. As dorsal hippocampus is more connected with spatial learning and memory, whereas the ventral region is related to emotional regulation, anxiety, and the stress response (Fanselow & Dong, 2010), we analyzed both regions separately. We first assessed acute cell proliferation and BDNF processing immediately following the adversity period at PND 13 (Fig. 2). SAM exposure did not alter acute cell proliferation, as indicated by similar densities of Ki67^+^ cells in both dorsal and ventral DG between groups.

**Figure 2.**
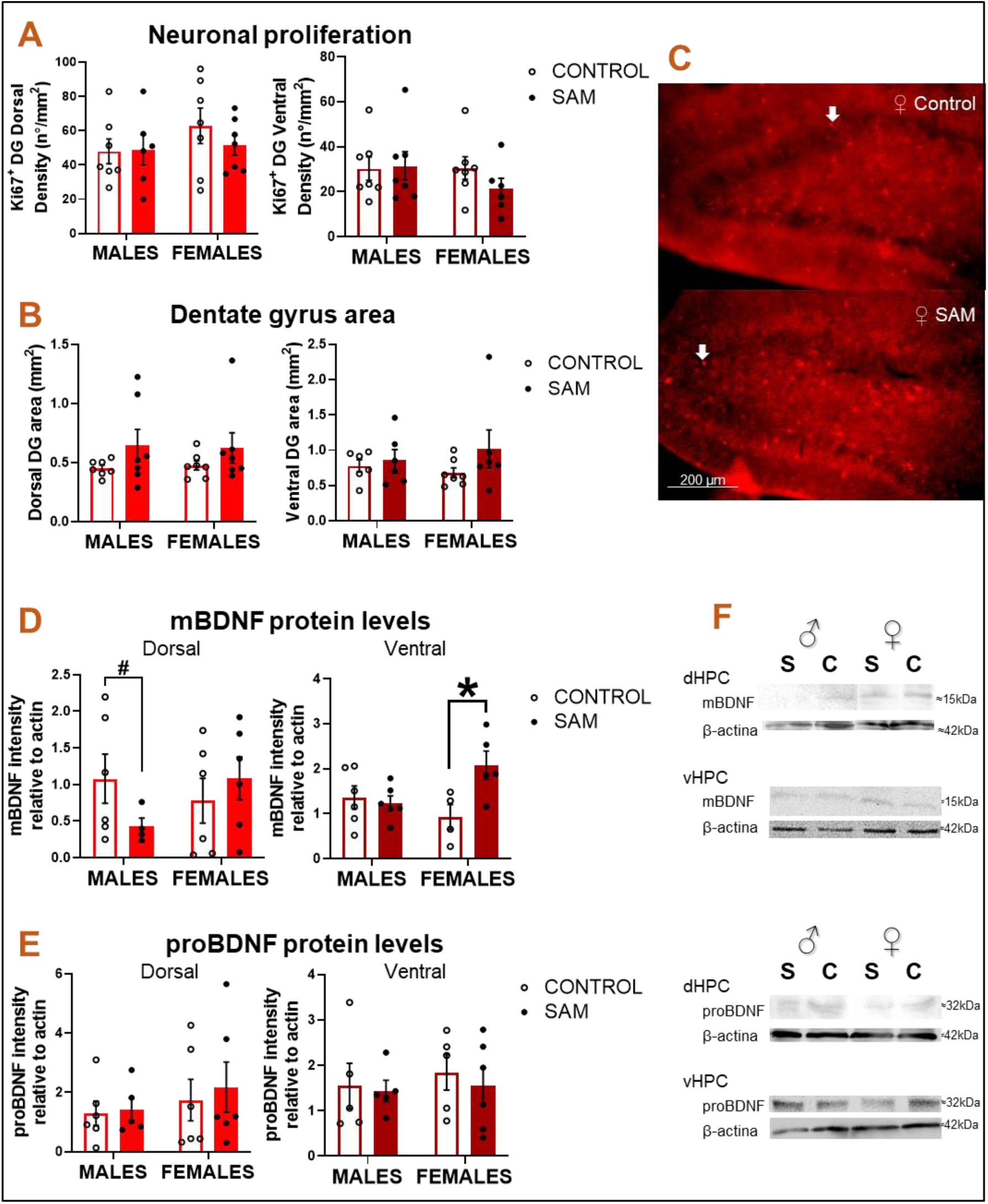
Hippocampal cell proliferation and BDNF protein expression at PND 13 following early-life adversity. (A) Quantification of Ki67^+^ cell density (cells/mm^2^) in dorsal and ventral DG. (B) Average area of the dorsal and ventral DG (mm^2^). (C) Representative micrographs of Ki67^+^ cells in the DG; scale bar = 200 µm. Arrows indicate representative Ki67^+^ positive nuclei. (D) Densitometric quantification of mBDNF and (E) pro-BDNF protein levels in dorsal and ventral hippocampus. (F) Representative Western blot bands for mature BDNF (mBDNF), pro-BDNF, and the loading control (β-actin) in the dorsal (dHPC) and ventral (vHPC) hippocampus. Bars represent mean ± SEM (n=6-8 per group in proliferation, and n=4-6 per group in Western blot). Two-way ANOVA followed by Tukey’s post-hoc test. * p < 0.05; # p < 0.10 (trend).

Western blot analysis of mature BDNF (mBDNF) and pro-BDNF revealed region- and sex-dependent effects at PND 13. SAM males exhibited a trend toward decreased mBDNF levels in the dorsal hippocampus (p=0.0988), whereas SAM females showed a significant upregulation of mBDNF in the ventral hippocampus compared to controls. Pro-BDNF levels remained unaltered across all conditions.

### Sex-dependent reduction in 1-week neuronal survival and ventral DG volume in juvenile SAM males

To examine whether early adversity affects the survival of newborn neurons, BrdU^+^ nuclei were quantified at PND 28 to assess 1-week-old cell survival (Fig. 3). SAM selectively impaired neuronal survival in males, evidenced by a significant reduction in BrdU^+^ 1-week-old cell density across both dorsal and ventral DG. In contrast, 1-week survival remained unaltered in SAM females. In addition, SAM males exhibited a trend toward decreased ventral DG area (p=0.0563).

**Figure 3.**
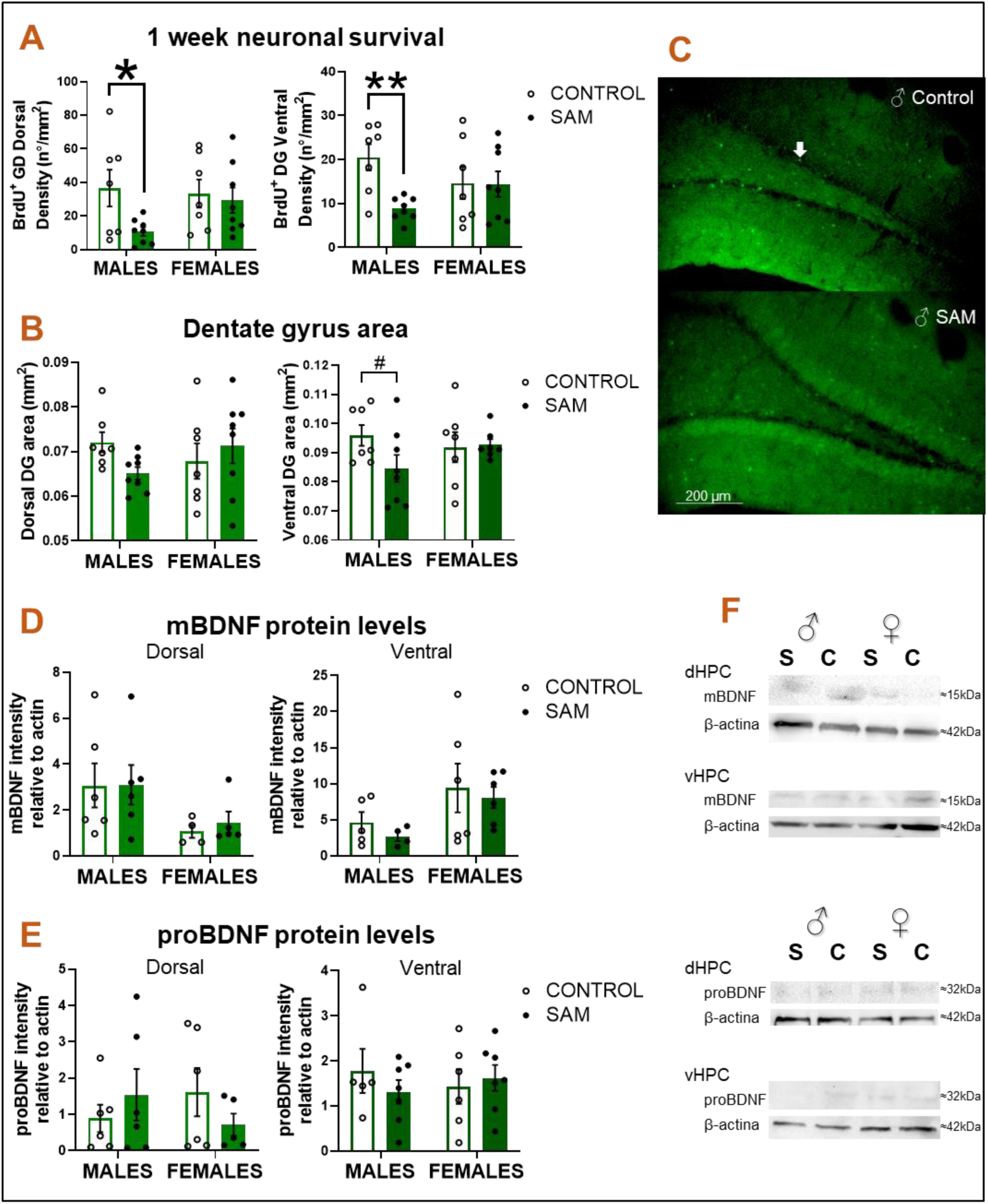
Hippocampal 1-week newborn cell survival and BDNF protein expression at PND 28 following early-life adversity. (A) Quantification of BrdU^+^ cell density (cells/mm^2^) in dorsal and ventral DG. (B) Average area of the dorsal and ventral DG (mm^2^). (C) Representative micrographs of BrdU^+^ cells in the DG; scale bar = 200 µm. Arrows indicate representative BrdU^+^ positive nuclei. (D) Densitometric quantification of mBDNF and (E) pro-BDNF protein levels in dorsal and ventral hippocampus. (F) Representative Western blot bands for mature BDNF (mBDNF), pro-BDNF, and the loading control (β-actin) in the dorsal (dHPC) and ventral (vHPC) hippocampus. Bars represent mean ± SEM (n=6-8 per group in neuronal survival analysis, and n=4-6 per group in Western blot). Two-way ANOVA followed by Tukey’s post-hoc test. ** p < 0.01; * p < 0.05; # p < 0.10 (trend).

At this developmental stage (PND 28), hippocampal mBDNF and pro-BDNF levels did not differ significantly between Control and SAM groups within each sex, although an overall main effect of sex on baseline mBDNF expression was detected in both hippocampal subregions (see Table 1).

## DISCUSSION

Early exposure to chronic stress disrupts the development and maturation of brain regions critical for emotional regulation, with consequences commonly evaluated in adulthood (Chen & Baram, 2016; Molet et al., 2014). However, behavioral alterations can emerge much earlier. Here, we demonstrate that early-life adversity induced by the SAM produces distinct juvenile behavioral phenotypes and sex-dependent alterations in hippocampal plasticity.

### Juvenile behavioral alterations

In line with previous findings in postnatal stress models (He et al., 2020; Molet et al., 2016), juvenile offspring did not display canonical anxiety-like behaviors. This absence is consistent with the notion that overt anxiety-related phenotypes typically consolidate in adulthood (Cui et al., 2020; Dalle Molle et al., 2012; Machado et al., 2013; Wang et al., 2012), as previously observed in adult SAM females (Grillo Balboa et al., 2025). Of note, SAM animals only showed a non-significant trend toward increased unsupported rearing in the open field test (p=0.06824), a subtle exploratory shift that may reflect early changes in emotional reactivity (Sturman et al., 2018).

In the forced swim test, juvenile SAM offspring showed increased immobility, corroborating reports that link resource-limitation models to passive coping strategies (Raineki et al., 2012; Rincón-Cortés & Sullivan, 2016; Roth; et al., 2013). Raineki et al. (2012) previously reported elevated immobility at PND 45 but not at PND 20 following SAM (PND 8–12). Our findings pinpoint PND 28 as the juvenile window where this passive coping phenotype emerges before persisting into adulthood (Rincón-Cortés & Sullivan, 2016). While the forced swim test was originally designed to assess antidepressant efficacy, it is often included in behavioral batteries to evaluate depressive-like features; however, we observed no anhedonic phenotype in the splash test. This highlights that increased immobility in juveniles may reflect a passive stress-coping strategy (Commons et al., 2017).

Spatial memory evaluation revealed deficits in the object location test (OLT) across both sexes, while novel object recognition (NOR) remained intact. Hippocampal dentate gyrus (DG) integrity is essential for spatial memory and pattern separation (Lajud & Torner, 2015). Early-life stress frequently impairs memory via neurogenic disruption, often showing male-biased vulnerability in adulthood (Naninck et al., 2015; Suri et al., 2013). However, Bath et al. (2017) demonstrated that while spatial memory deficits persist into adulthood only in males, females display a transient impairment during the juvenile stage. Our results confirm that the juvenile period represents a critical window where spatial deficits manifest across both sexes, although likely through diverging mechanisms. In males, OLT deficits aligned with reduced 1-week neuronal survival. In females, memory impairment occurred independently of this survival marker, suggesting that other neurodevelopmental factors - such as dendritic complexity or synaptic plasticity (Nicolas et al., 2022) - drive the behavioral phenotype. Moreover, because 1-week-old neurons are still immature, future assessments of fully differentiated, mature adult-born neurons will be necessary to further elucidate neurogenesis-dependent cognitive performance.

### Sex-dependent deficits in hippocampal neurogenesis

SAM altered hippocampal neurogenesis in a sex-dependent manner, reducing 1-week-old neuronal survival across both dorsal and ventral DG exclusively in males. While literature in adolescent rodents is sparse, previous reports show reduced proliferation in stressed adolescent males (Liu et al., 2026) and preserved neurogenesis alongside dendritic retraction in juvenile females (Nicolas et al., 2022). In adulthood, postnatal stress robustly impairs hippocampal neuronal survival and cognition in males, while sparing or exerting attenuated effects in females (Loi et al., 2014; Naninck et al., 2015; Oomen et al., 2009; Suri et al., 2013; Zuena et al., 2008).

This sexual divergence stems from early perinatal brain organization (McEwen, 2017) and endocrine events such as mini-puberty (PND 9–20 in rats) (Devillers et al., 2022). Because estrogens exert neuroprotective effects and regulate the balance between neural stem/progenitor cell proliferation and differentiation (Bustamante-barrientos et al., 2021), the transient estrogen surge during female mini-puberty may confer neuroprotection against SAM-induced cell loss, a hypothesis interesting to challenge in the future.

Paralleling clinical evidence where early neglect correlates with reduced hippocampal volume preferentially in males (Frodl et al., 2010), we observed a trend toward reduced ventral DG volume in SAM males (p=0.0563). This structural reduction is biologically consistent with the impaired neuronal survival observed in this group and mirrors findings by Naninck et al. (2015) at PND 9 and adulthood. Notably, previous data from our laboratory demonstrated that adult SAM males but not females, develop anhedonia and motivational deficits (Grillo Balboa et al., 2025). The selective reduction in juvenile male neuronal survival reported here may represent the early biological substrate governing this sex-divergent trajectory.

### Early sex-specific BDNF regulation

Neurotrophins act as key regulators of this neurogenic process (Wu & Zhang, 2023). Among them, mature BDNF (mBDNF) promotes neuronal differentiation, survival, and synaptic plasticity. Conversely, its precursor, proBDNF, exerts opposing, pro-apoptotic effects through distinct receptor signaling pathways, such that the dynamic balance between mBDNF and proBDNF levels critically dictates the rate and outcome of hippocampal neurogenesis (Foltran & Diaz, 2016).

In this work, at PND 13, SAM induced a significant increase in mBDNF levels in the ventral hippocampus of females without altering proBDNF levels, effectively shifting the balance toward pro-survival signaling. This contrasts with a trend toward reduced mBDNF levels in the male dorsal hippocampus (p=0.0988), an early divergence that aligns with the subsequent sex-specific differences observed in juvenile neuronal survival.

The functional crosstalk between sex steroids and BDNF signaling is well established, as estrogens can positively regulate BDNF expression and downstream TrkB activation (Chan & Ye, 2017). In this context, alongside known sex differences in dentate gyrus mBDNF levels during the second postnatal week (Sardar et al., 2021), the possibility exists that the transient estrogen surge occurring during female mini-puberty (Devillers et al., 2022) may drive a compensatory upregulation of ventral mBDNF in response to SAM, potentially buffering the neurogenic niche against stress-induced cell loss. The absence of changes in acute cell proliferation at PND 13 indicates that early shifts in BDNF signaling may primarily prime postmitotic neuronal differentiation and survival rather than initial cell division. In addition, other signaling pathways - such as glucocorticoid receptor activation, neuroinflammatory cascades, or Wnt signaling - could be acting in parallel to modulate hippocampal neurogenesis in this model (El-Kadi et al., 2024; Ortega-martínez, 2015).

## REFERENCES

Agorastos, A., Pervanidou, P., Chrousos, G. P., & Baker, D. G. (2019). Developmental trajectories of early life stress and trauma: A narrative review on neurobiological aspects beyond stress system dysregulation. Frontiers in Psychiatry, 10(MAR), 1–25. 10.3389/fpsyt.2019.00118

Apple, D. M., Fonseca, R. S., & Kokovay, E. (2016). The role of adult neurogenesis in psychiatric and cognitive disorders. Brain Research, 1–9. 10.1016/j.brainres.2016.01.023

Bath, K. G., Goodwill, H., Sciences, P., States, U., & States, U. (2017). Early life stress accelerates behavioral and neural maturation of the hippocampus in male mice. Horm Behav, 82, 64–71. 10.1016/j.yhbeh.2016.04.010.Early

Bustamante-barrientos, F. A., Méndez-ruette, M., Ortloff, A., Luz-crawford, P., Rivera, F. J., Figueroa, C. D., Molina, L., & Morales-garcia, J. A. (2021). The Impact of Estrogen and Estrogen-Like Molecules in Neurogenesis and Neurodegeneration : Beneficial or Harmful? 15(March), 1–19. 10.3389/fncel.2021.636176

Callaghan, B., Meyer, H., Opendak, M., Van Tieghem, M., Harmon, C., Li, A., Lee, F. S., Sullivan, R. M., & Tottenham, N. (2019). Using a Developmental Ecology Framework to Align Fear Neurobiology Across Species. Annual Review of Clinical Psychology, 15, 345–369. 10.1146/annurev-clinpsy-050718-095727

Chan, C. B., & Ye, K. (2017). Sex Differences in Brain-Derived Neurotrophic Factor Signaling and Functions. J Neurosci Res., 95, 328–335. 10.1002/jnr.23863.Sex

Chen, W., Zhang, Q., Su, W., Zhang, H., Yang, Y., Qiao, J., Sui, N., & Li, M. (2014). Effects of 5-hydroxytryptamine 2C receptor agonist MK212 and 2A receptor antagonist MDL100907 on maternal behavior in postpartum female rats. Pharmacology Biochemistry and Behavior, 117, 25–33. 10.1016/j.pbb.2013.11.034

Chen, Y., & Baram, T. Z. (2016). Toward understanding how early-life stress reprograms cognitive and emotional brain networks. Neuropsychopharmacology, 41(1), 197–206. 10.1038/npp.2015.181

Colapietro, A. A., Grillo Balboa, J., Ceol Retamal, M. N., Regueira, E., Hermida, G. N., Cantarelli, V. I., Ponzio, M. F., Pallarés, M. E., Antonelli, M. C., & Diaz, S. L. (2025). Infant Maltreatment Induces Early Alterations in Adrenal Glands and Stress Response in Juvenile Rat Offspring. Neurochemical Research, 50:108. 10.1007/s11064-025-04363-5

Commons, K. G., Cholanians, A. B., Babb, J. A., & Ehlinger, D. G. (2017). The Rodent Forced Swim Test Measures Stress-Coping Strategy, Not Depression-like Behavior. ACS Chemical Neuroscience, 8(5), 955–960. 10.1021/acschemneuro.7b00042

Condon, E., Sadler, L., & Mayes, L. (2018). Toxic stress and protective factors in multi-ethnic school age children: A research protocol. Research in Nursing and Health, 1, 8. 10.1002/nur.21851

Cui, Y., Cao, K., Lin, H., Cui, S., Shen, C., Wen, W., Mo, H., Dong, Z., Bai, S., Yang, L., Shi, Y., & Zhang, R. (2020). Early-Life Stress Induces Depression-Like Behavior and Synaptic-Plasticity Changes in a Maternal Separation Rat Model: Gender Difference and Metabolomics Study. Frontiers in Pharmacology, 11(February), 1–13. 10.3389/fphar.2020.00102

Dalle Molle, R., Portella, A. K., Goldani, M. Z., Kapczinski, F. P., Leistner-Segala, S., Salum, G. A., Manfro, G. G., & Silveira, P. P. (2012). Associations between parenting behavior and anxiety in a rodent model and a clinical sample: Relationship to peripheral BDNF levels. Translational Psychiatry, 2(11), e195–8. 10.1038/tp.2012.126

Devillers, M. M., Mhaouty-kodja, S., & Guigon, C. J. (2022). Deciphering the Roles & Regulation of Estradiol Signaling during Female Mini-Puberty : Insights from Mouse Models. International Journal of Molecular Sciences, 23(13695). doi10.3390/ijms232213695

El-Kadi, R. A., AbdelKader, N. F., Zaki, H. F., & Kamel, A. S. (2024). Influence of β-catenin signaling on neurogenesis in neuropsychiatric disorders: Anxiety and depression. Drug Dev Res., 85(e22157). doi: 10.1002/ddr.22157

Fanselow, M. S., & Dong, H.-W. (2010). Are The Dorsal and Ventral Hippocampus functionally distinct structures*?* 65(1), 1–25. 10.1016/j.neuron.2009.11.031.

Felitti, V., Anda, R., & Nordenberg, D. (1998). Relationship of childhood abuse and household dysfunction to many of the leading causes of death in adults: The Adverse Childhood Experiences (ACE) Study. American Journal of Preventive Medicine., 14(4), 245–258. 10.1016/S0749-3797(98)00017-8

Foltran, R. B., & Diaz, S. L. (2016). BDNF isoforms: a round trip ticket between neurogenesis and serotonin? Journal of Neurochemistry, 204–221. 10.1111/jnc.13658

Frodl, T., Reinhold, E., Koutsouleris, N., Reiser, M., & Meisenzahl, E. M. (2010). Interaction of childhood stress with hippocampus and prefrontal cortex volume reduction in major depression. Journal of Psychiatric Research, 44(13), 799–807. 10.1016/j.jpsychires.2010.01.006

Gage, F. H. (2000). Mammalian Neural Stem Cells. Science, 287(1433). 10.1126/science.287.5457.1433 This

Grillo Balboa, J., Colapietro, A. A., Cantarelli, V. I., Ponzio, M. F., Retamal, M. N. C., Pallarés, M. E., Antonelli, M. C., & Chertoff, M. (2025). Sex-Specific Outcomes in a Rat Model of Early-Life Stress Due to Adverse Caregiving. Neurotoxicity Research, 43(10), 1–18. 10.1007/s12640-025-00731-9

Hattiangady, B., Mishra, V., Kodali, M., Shuai, B., Rao, X., & Shetty, A. K. (2014). Object location and object recognition memory impairments, motivation deficits and depression in a model of Gulf War illness. Frontiers in Behavioral Neuroscience, 8(MAR), 1–10. 10.3389/fnbeh.2014.00078

He, T., Guo, C., Wang, C., Hu, C., & Chen, H. (2020). Effect of early life stress on anxiety and depressive behaviors in adolescent mice. Brain and Behavior, 10(3), 1–10. 10.1002/brb3.1526

Heim, C., & Nemeroff, C. B. (2001). The Role of Childhood Trauma in the Neurobiology of Mood and Anxiety Disorders : Preclinical and Clinical Studies.

Kempermann, G., Song, H., & Gage, F. H. (2015). Neurogenesis in the Adult Hippocampus. 1–14.

Lajud, N., & Torner, L. (2015). Early life stress and hippocampal neurogenesis in the neonate: Sexual dimorphism, long term consequences and possible mediators. Frontiers in Molecular Neuroscience, 8(FEB), 1–10. 10.3389/fnmol.2015.00003

Liu, Z., Wang, J., Ge, Y., Wang, Y., Ling, H., Liu, Y., Zhang, J., You, Z., & Han, Y. (2026). Brain Behavior and Immunity PPAR γ in microglia helps protect adolescent male mice from harmful effects of stress during early development. Brain Behavior and Immunity, 134(February), 106483. 10.1016/j.bbi.2026.106483

Loi, M., Koricka, S., Lucassen, P. J., & Joëls, M. (2014). Age- and sex-dependent effects of early life stress on hippocampal neurogenesis. 5(February), 1–12. 10.3389/fendo.2014.00013

Machado, T. D., Dalle Molle, R., Laureano, D. P., Portella, A. K., Werlang, I. C. R., Benetti, C. D. S., Noschang, C., & Silveira, P. P. (2013). Early life stress is associated with anxiety, increased stress responsivity and preference for “comfort foods” in adult female rats. Stress, 16(5), 549–556. 10.3109/10253890.2013.816841

McEwen, B. S. (2017). Neurobiological and Systemic Effects of Chronic Stress. Chronic Stress, 1. 10.1177/2470547017692328

Molet, J., Heins, K., Zhuo, X., Mei, Y. T., Regev, L., Baram, T. Z., & Stern, H. (2016). Fragmentation and high entropy of neonatal experience predict adolescent emotional outcome. Translational Psychiatry, 6(702). 10.1038/tp.2015.200

Molet, Jenny, Maras, P. M., Avishai-Eliner, S., & Baram, T. Z. (2014). Naturalistic rodent models of chronic early-life stress. Developmental Psychobiology, 56(8), 1675–1688. 10.1002/dev.21230

Moriceau, S., Shionoya, K., Jakubs, K., & Sullivan, R. M. (2009). Early-Life Stress Disrupts Attachment Learning: The Role of Amygdala Corticosterone, Locus Ceruleus Corticotropin Releasing Hormone, and Olfactory Bulb Norepinephrine. The Journal of Neuroscience, 29(50)(15745), 15755. 10.1523/JNEUROSCI.4106-09.2009

Naninck, E. F. G., Hoeijmakers, L., Kakava-Georgiadou, N., Meesters, A., Lazic, S. E., Lucassen, P. J., & Korosi, A. (2015). Chronic early life stress alters developmental and adult neurogenesis and impairs cognitive function in mice. Hippocampus, 25(3), 309–328. 10.1002/hipo.22374

Nicolas, S., McGovern, A. J., Hueston, C. M., O’Mahony, S. M., Cryan, J. F., O’Leary, O. F., & Nolan, Y. M. (2022). Prior maternal separation stress alters the dendritic complexity of new hippocampal neurons and neuroinflammation in response to an inflammatory stressor in juvenile female rats. Brain Behav Immun, 99, 327–338. doi: 10.1016/j.bbi.2021.10.016

Oomen, C. A., Girardi, C. E. N., Cahyadi, R., Verbeek, E. C., Krugers, H., Joe, M., & Lucassen, P. J. (2009). Opposite Effects of Early Maternal Deprivation on Neurogenesis in Male versus Female Rats. 4(1). 10.1371/journal.pone.0003675

Ortega-martínez, S. (2015). Influences of prenatal and postnatal stress on adult hippocampal neurogenesis : The double neurogenic niche hypothesis. Behavioural Brain Research, 281, 309–317. 10.1016/j.bbr.2014.12.036

Pallarés, M. E., Monteleone, M. C., Pastor, V., Grillo Balboa, J., Alzamendi, A., Brocco, M. A., & Antonelli, M. C. (2021). Early-Life Stress Reprograms Stress-Coping Abilities in Male and Female Juvenile Rats. Molecular Neurobiology, 58(11), 5837–5856. 10.1007/s12035-021-02527-2

Pastor, V., Pallarés, M. E., & Antonelli, M. C. (2018). Prenatal stress increases adult vulnerability to cocaine reward without affecting pubertal anxiety or novelty response. Behavioural Brain Research, 339, 186–194. 10.1016/j.bbr.2017.11.035

Raineki, C., Cortés, M., Belnoue, L., & Sullivan, R. (2012). Effects of early-life abuse differ across development: Infant social behavior deficits are followed by adolescent depressive-like behaviors mediated by the Amygdala. Journal of Neuroscience, 32(22), 7758–7765. 10.1523/JNEUROSCI.5843-11.2012

Raineki, C., Moriceau, S., & Sullivan, R. (2010). Developing a Neurobehavioral Animal Model of Infant Attachment to an Abusive Caregiver. Biological Psychiatry, 67(12), 1137–1145. 10.1016/j.biopsych.2009.12.019

Rincón-Cortés, M., & Sullivan, R. M. (2016). Emergence of social behavior deficit, blunted corticolimbic activity and adult depression-like behavior in a rodent model of maternal maltreatment. Translational Psychiatry, 6, 930. 10.1038/tp.2016.205

Roth;, T. L., Raineki;, C., Salstein;, L., Perry;, R., Wilson;, T. A. S., Sloan;, A., Lalji;, B., Hammock;, E., Wilson;, D. A., Levitt;, P., Okutani;, F., Kaba;, H., & Sullivan, R. M. (2013). Neurobiology of secure infant attachment and attachment despite adversity: a mouse model. Early Human Development, 12(7), 673–680. 10.1016/j.earlhumdev.2006.05.022

Roth, T. L., & Sullivan, R. M. (2005). Memory of early maltreatment: Neonatal behavioral and neural correlates of maternal maltreatment within the context of classical conditioning. Biological Psychiatry, 57(8), 823–831. 10.1016/j.biopsych.2005.01.032

Roversi, K., de David Antoniazzi, C. T., Milanesi, L. H., Rosa, H. Z., Kronbauer, M., Rossato, D. R., Duarte, T., Duarte, M. M., & Burger, M. E. (2019). Tactile Stimulation on Adulthood Modifies the HPA Axis, Neurotrophic Factors, and GFAP Signaling Reverting Depression-Like Behavior in Female Rats. Molecular Neurobiology, 56(9), 6239–6250. 10.1007/s12035-019-1522-5

Sardar, R., Zandieh, Z., Namjoo, Z., Soleimani, M., Shirazi, R., & Hami, J. (2021). Laterality and sex differences in the expression of brain-derived neurotrophic factor in developing rat hippocampus. Metabolic Brain Disease, 36, 133–144. 10.1007/s11011-020-00620-4

Sturman, O., Germain, P. L., & Bohacek, J. (2018). Exploratory rearing: a context- and stress-sensitive behavior recorded in the open-field test. Stress, 21(5), 443–452. 10.1080/10253890.2018.1438405

Suri, D., Veenit, V., Sarkar, A., Thiagarajan, D., Kumar, A., Nestler, E. J., Galande, S., & Vaidya, V. A. (2013). Early stress evokes age-dependent biphasic changes in hippocampal neurogenesis, Bdnf expression, and cognition. Biological Psychiatry, 73(7), 658–666. 10.1016/j.biopsych.2012.10.023

Wang, X. D., Labermaier, C., Holsboer, F., Wurst, W., Deussing, J. M., Müller, M. B., & Schmidt, M. V. (2012). Early-life stress-induced anxiety-related behavior in adult mice partially requires forebrain corticotropin-releasing hormone receptor 1. European Journal of Neuroscience, 36(3), 2360–2367. 10.1111/j.1460-9568.2012.08148.x

Wu, A., & Zhang, J. (2023). Neuroinflammation, memory, and depression : new approaches to hippocampal neurogenesis. Journal of Neuroinflammation, 1–20. 10.1186/s12974-023-02964-x

Zuena, A. R., Mairesse, J., Casolini, P., Cinque, C., Alema, G. S., Morley-fletcher, S., Chiodi, V., Spagnoli, L. G., Gradini, R., Catalani, A., Nicoletti, F., & Maccari, S. (2008). Prenatal Restraint Stress Generates Two Distinct Behavioral and Neurochemical Profiles in Male and Female Rats. 3(5). 10.1371/journal.pone.0002170

